# Toxic PR poly-dipeptides encoded by the *C9orf72* repeat expansion target Kapβ2 and dysregulate phase separation of low-complexity domains

**DOI:** 10.1101/812099

**Authors:** Hitoki Nanaura, Honoka Kawamukai, Ayano Fujiwara, Takeru Uehara, Mari Nakanishi, Tomo Shiota, Masaki Hibino, Yuichiro Aiba, Pattama Wiriyasermkul, Sotaro Kikuchi, Riko Nagata, Masaya Matsubayashi, Shushi Nagamori, Osami Shoji, Koichiro Ishimori, Hiroyoshi Matsumura, Kazuma Sugie, Tomohide Saio, Takuya Yoshizawa, Eiichiro Mori

**Author notes:** co-first (Hitoki Nanaura, Honoka Kawamukai). To whom correspondence may be addressed: Department of Chemistry, Faculty of Science, Hokkaido University, Sapporo, Hokkaido 060-0810, Japan. To whom correspondence may be addressed: College of Life Sciences, Ritsumeikan University, Kusatsu, Shiga 525-8577, Japan. To whom correspondence may be addressed: Department of Future Basic Medicine, Nara Medical University, Kashihara, Nara 634-8521, Japan.

## Abstract

Low-complexity (LC) domains of proteins are found in about one fifth of human proteome, and a group of LC-domains form labile cross-β polymers and liquid-like droplets. Polymers and droplets formed from LC-domains are dynamically regulated by posttranslational modifications and molecular chaperones including nuclear transport receptors. Repeat expansion in the first intron of a gene designated *C9orf72*, which is the most prevalent form of familial amyotrophic lateral sclerosis (ALS), causes nucleocytoplasmic transport deficit, however, the detailed mechanism remains unsolved. Here we show that the proline:arginine (PR) poly-dipeptides encoded by the *C9orf72* repeat expansion bound nuclear transport receptor Kapβ2 through its nuclear localization signal (NLS) recognition motif, and inhibited the ability of Kapβ2 to melt fused in sarcoma (FUS) droplets by competing interaction with FUS. The findings in this study offer mechanistic insights as to how the *C9orf72* repeat expansion disables nucleocytoplasmic transport and causes neurodegenerative diseases.

---

Intrinsically disordered low-complexity (LC) protein sequences (LC-domains), also called prion-like domains or amyloid-like domains, are often found in RNA/DNA-binding proteins, intermediate filaments and nuclear pore proteins (1–3). Approximately 20% of human proteome harbors LC-domains, and a group of LC-domains forms labile cross-β polymers and liquid-like droplets in the test tube and RNA granules in the cell (1,4). RNA-binding proteins fused in sarcoma (FUS) and heterogeneous nuclear ribonucleo-proteins (hnRNPs) have been extensively studied in recent years, including studies on their capacity to phase-separate into liquid-like droplets both in the test tube and in the cell (1,4–7).

Labile cross-β polymers formed from LC-domains (hereafter LC polymers) are controlled to be dynamic and not to grow too long at the physiological condition in the cell (8). Once the balance between short and long tilts toward long, LC polymers turn into stable pathogenic fibrils, which are often found in neurodegenerative diseases (8). Human genetics has provided us with some clues to how LC polymers become pathogenic. Specific mutations [D-to-V] in the LC domain of hnRNPs from amyotrophic lateral sclerosis (ALS) and multisystem proteinopathy (MSP) cause stable cross-β polymers (2,9). A hexanucleotide repeat expansion in *C9orf72*, the most prevalent form of familial ALS and frontotemporal dementia (FTD), produces the most toxic products–poly-dipeptides, proline:arginine (PR) and glycine:arginine (GR) (10,11). These arginine-rich poly-dipeptides bind proteins with LC sequences (2,12) and stabilize LC polymers (2).

Cells have multiple layers of regulatory system to keep LC polymers short and dynamic. Most posttranslational modifications (e.g. phosphorylation, oxidation) are found at LC regions and well known to regulate LC polymerization (1,5–7,10,13–15). Recent works by four different groups similarly conclude that nuclear transport receptors function as a molecular chaperone to keep the LC proteins monomeric (14,16–18).

Three genetic studies revealed the repeat-expansion in *C9orf72* gene disrupts nucleocytoplasmic transport (19–21). One common path for both nuclear import and export is the pore buried in the nuclear envelop called nuclear pores. PR poly-dipeptides derived from the *C9orf72* repeat expansion bind and stabilize polymeric form, but not monomeric form, of phenylalanine:glycine (FG)-rich domains of nuclear pore proteins (3). Nuclear transport receptors are known to be responsible for nucleocytoplasmic transport, but it is still unclear how the repeat expansion in *C9orf72* compromises nucleocytoplasmic transport.

Although the previous studies have reported that PR poly-dipeptides bind and stabilize the cross-β polymer conformation of the LC proteins, the effect of the PR poly-dipeptides to the regulators including nuclear transporter is remained elusive. In this study, we have investigated the interaction between PR poly-dipeptides and nuclear transport receptor Kapβ2 and evaluated the effect of the PR poly-dipeptides to the function of Kapβ2 in regulation of the liquid-liquid phase separation of LC proteins. Our data from in vitro experiments exploiting biophysical methods provide a mechanistic insight into as to how PR poly-dipeptides disable nucleocytoplasmic transport and cause neurodegenerative diseases.

## Results

### PR poly-dipeptides inhibit chaperone function of Kapβ2

First, we examined the effect of PR poly-dipeptides on Kapβ2 in terms of melting phase-separated full-length FUS droplets. We prepared 18 repeated proline:arginine poly-dipeptide with maltose binding protein (MBP) motif (MBP-PR18). As previously reported (16), FUS forms liquid-like droplets via liquid-liquid phase separation, and the phase separation of FUS is blocked by Kapβ2 observed with fluorescence microscopy (Fig. 1A). In contrast, Kapβ2 loses the ability to melt the FUS droplet in the presence of MBP-PR18 (Fig. 1A). To better clarify the data, we performed further analysis on the formation and dissolution of FUS droplet by monitoring the turbidity of the solution. Phase-separated droplets floating in the solution demonstrate turbidity and the progress of phase separation can be monitored by tracking turbidity. Full length of FUS formed liquid droplets, and Kapβ2 blocked FUS from phase separating into liquid-like droplets. By contrast, the addition of equimolar of MBP-PR18 inhibited the ability of Kapβ2 (Fig. 1B). These data indicate that PR poly-dipeptides with Kapβ2 inhibited the chaperone function of Kapβ2 on FUS phase separation.

**Figure 1.**
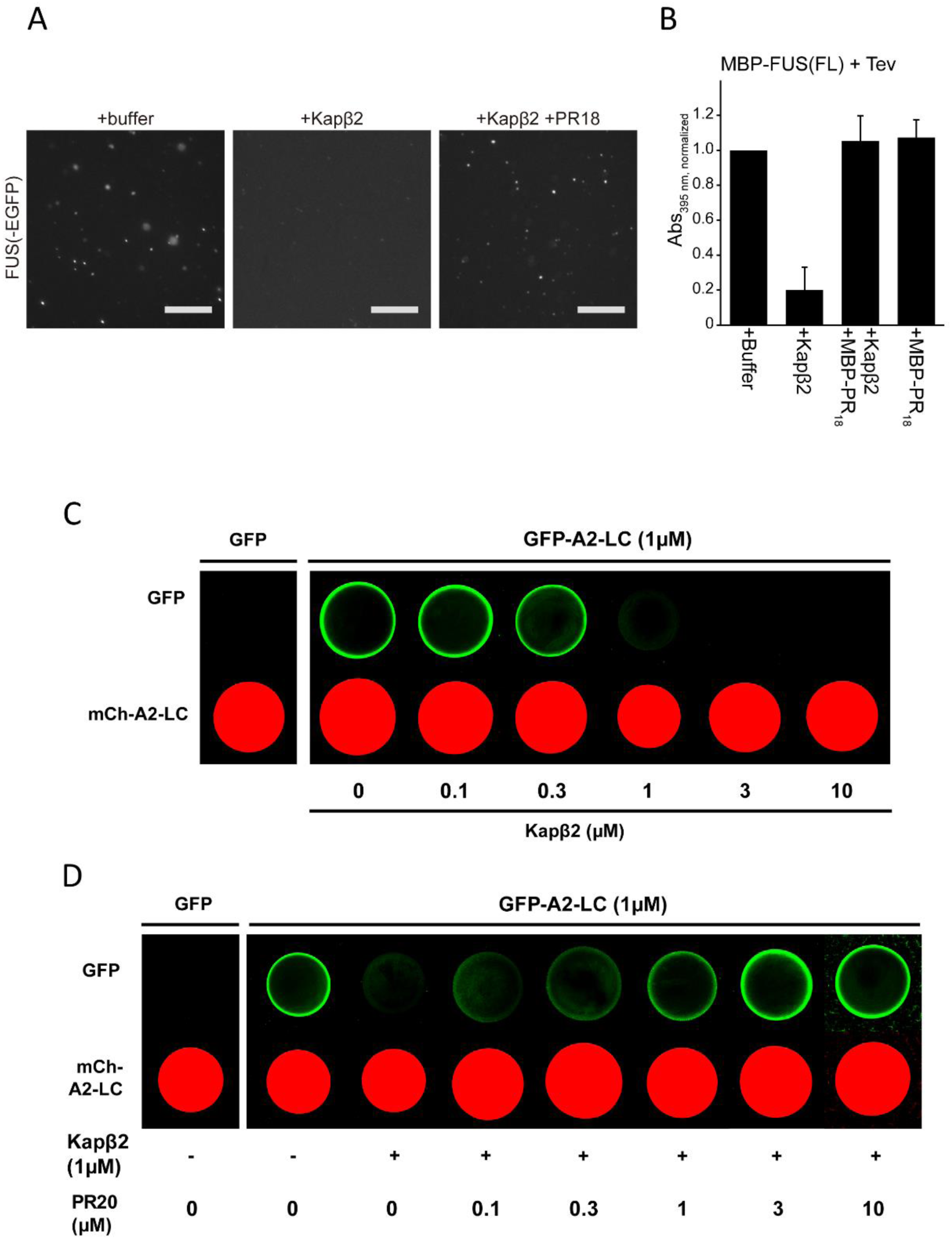
PR poly-dipeptides inhibit chaperone function of Kapβ2. (A) Mixture of 7.2μM MBP-FUS and 0.8μM MBP-FUS-EGFP were treated with TEV for 1 hour in presence or absence of Kapβ2 and MBP-PR18. 50μm scale bars are shown in the images. (B) Turbidity of 8μM MBP-FUS in the presence of buffer, ± 8μM Kapβ2, ± 8μM MBP-PR18. Abs395 nm is normalized to measurement of MBP-FUS + buffer + Tev. Mean of 3 technical replicates, ±SD. (C) Hydrogel droplets made from mCherry:LC domain of hnRNPA2 (lower images) were incubated with 1µM of GFP (left panel) and GFP:hnRNPA2-LC (right panel) and visualized by confocal microscopy. GFP:hnRNPA2-LC was challenged for homotypic polymer extension in the presence of different concentration of Kapβ2 (left to right: 0.1µM, 0.3µM, 1µM, 3µM, 10µM, respectively). (D) GFP:hnRNPA2-LC containing Kapβ2 (1.0µM) were applied to hydrogel droplets in the presence of diverse concentration of PR20-HA (left to right: 0.1µM, 0.3µM, 1µM, 3µM, 10µM, respectively).

Next, we prepared the LC domain of FUS (residues 2-214), and tested the ability of Kapβ2 in preventing polymerization of the LC domain of FUS using hydrogel binding assay (1). Briefly, the FUS-LC fused to mCherry was purified as described previously (1), and concentrated to self-associate and form cross-β polymers to obtain hydrogel droplets. After that the prepared GFP:FUS-LC was challenged for homotypic polymer extension in the presence of Kapβ2 (see more details in Methods). FUS-LC hydrogel binding was not blocked by Kapβ2 (Fig. S1A), suggesting the importance of proline-tyrosine nuclear localization signal (PY-NLS) for Kapβ2 to recognize FUS as a substrate.

Then, we applied the LC domain of hnRNPA2 (residues 181-341), whose PY-NLS is located in the middle of its LC domain. Kapβ2 (1μM) was capable of blocking hydrogel binding of GFP:hnRNPA2-LC (1μM) (Fig. 1C), suggesting that Kapβ2 captures hnRNPA2-LC in 1:1 ratio by the interaction between PY-NLS and monomeric state of hnRNPA2-LC. Further, 1μM of PR poly-dipeptides inhibited the ability of Kapβ2 from blocking the hydrogel binding of hnRNPA2-LC (Fig. 1D).

### Interaction between Kapβ2 and PR poly-dipeptides

Based on two independent interactomes, Karyopherins are found to be potential interactors in the complex with PR poly-dipeptides (2,12). To see the direct interaction between PR poly-dipeptides and Kapβ2, we performed on-column binding assay with purified recombinant proteins (Fig. 2A, B). We found that both GST-PR18 and MBP-PR18 bound to Kapβ2 by the on-column binding assay, while PR8 (eight repeat poly-dipeptide of Pro-Arg) did not bind (Fig. 2A, B). Isothermal titration calorimetry (ITC) showed that MBP-PR18 binds to Kapβ2 in 1:1 ratio at *K*_d_ value of 81.3nM (Fig. 2C). The binding assay indicates that Kapβ2 interacts with PR poly-dipeptides at one site with high affinity. Kapβ2 binds PR poly-dipeptides in a similar affinity to PY-NLS of FUS and stronger than full-length FUS (Table 1, Table S1). These results indicate that expanded PR poly-dipeptides directly interact and disrupt Kapβ2 to inhibit droplet formation of FUS.

**Figure 2.**
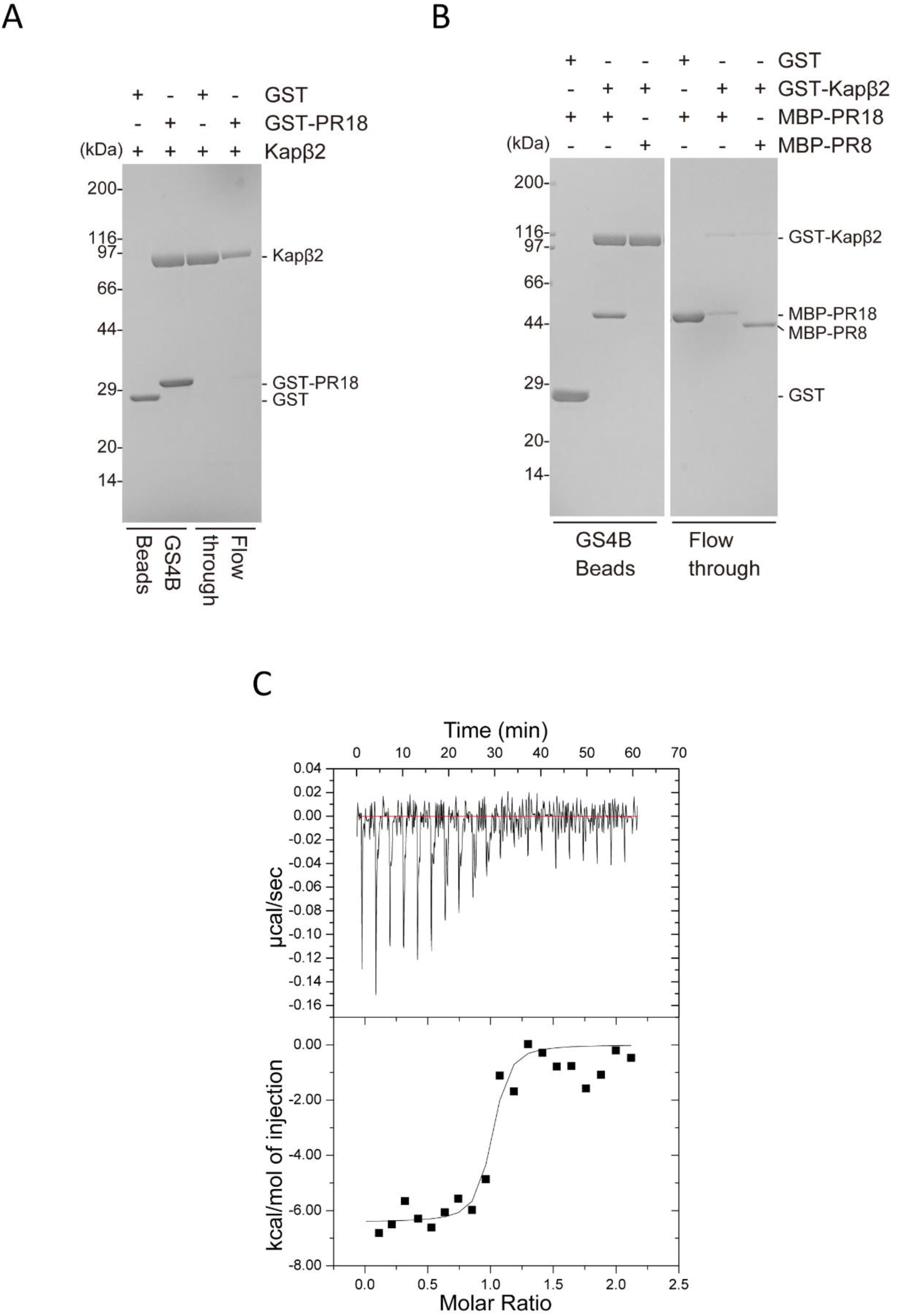
Interaction between Kapβ2 and PR poly-dipeptides. (A) Pull-down binding assay showing interaction between GST-PR18 and Kapβ2. (B) Pull-down binding assay showing interaction between GST-Kapβ2 and MBP-PR18/MBP-PR8. (C) Dissociation constants (Kd) measured by ITC of Kapβ2(ΔLoop) binding to MBP-PR18.

**Table 1.**
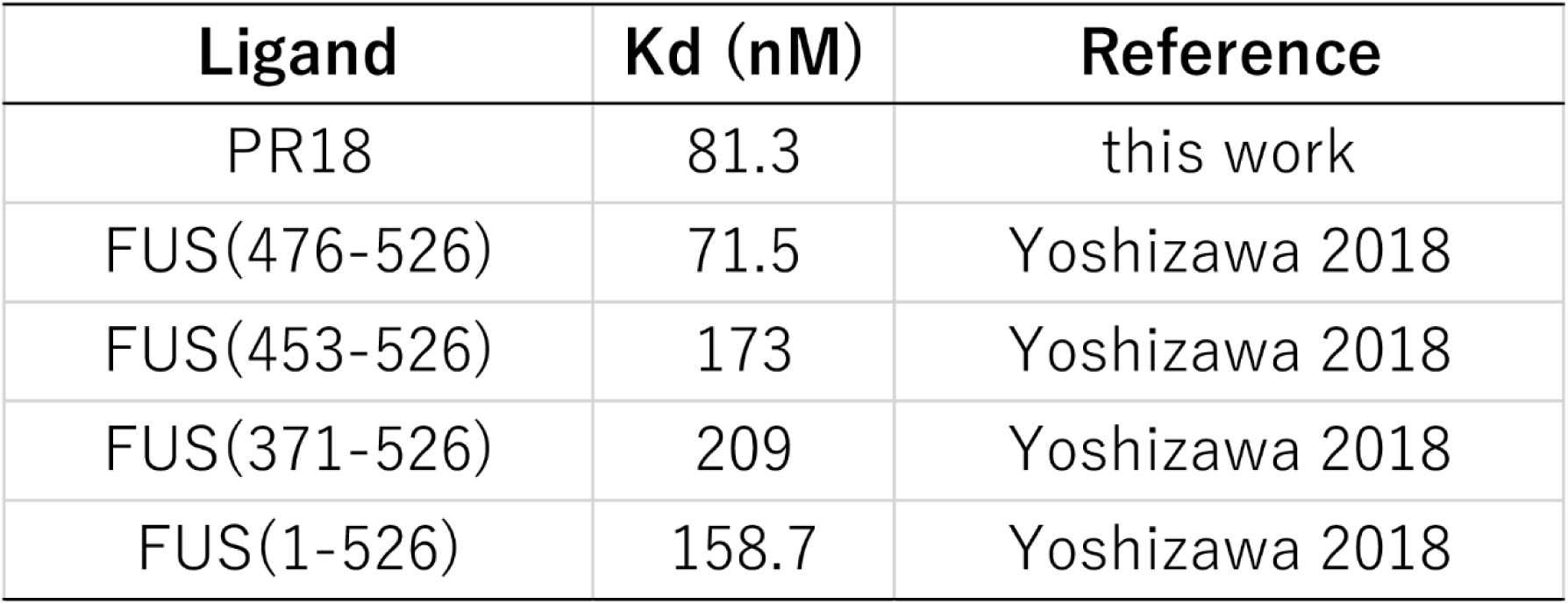
Kd value of MBP-tagged of PR18 and FUS fragments against Kapβ2 determined by ITC

| **Ligand** | **K<sub>d</sub> (nM)** | **Reference** |
| --- | --- | --- |
| PR18 | 81.3 | this work |
| FUS(476-526) | 71.5 | Yoshizawa 2018 |
| FUS(453-526) | 173 | Yoshizawa 2018 |
| FUS(371-526) | 209 | Yoshizawa 2018 |
| FUS(1-526) | 158.7 | Yoshizawa 2018 |

### Interaction between Kapβ2 and PR poly-dipeptides investigated by solution NMR

The presence of a direct interaction between Kapβ2 and PR poly-dipeptides was further validated by solution nuclear magnetic resonance (NMR) (Fig. 3). Despite large size of Kapβ2 (100 kDa) in the solution NMR, the use of advanced NMR techniques including methyl-selective isotope labeling (Fig. 3A) as well as methyl-transverse relaxation-optimized spectroscopy (TROSY) (22,23), achieved high-quality NMR spectra (Fig. 3B). Although the NMR resonances are only derived from the methyl groups of Ile, Leu, Val, and Met, these methyl groups are found to be widely distributed in the structure of Kapβ2 and thus, expected to sensitively detect the binding of poly-dipeptides to any region in Kapβ2. The NMR spectra of the isotopically labeled Kapβ2 were acquired in the absence and presence of PR20 (Fig. 3C). The addition of the PR poly-dipeptides induced significant perturbations to several resonances of Kapβ2 (Fig. 3C), indicating that the PR poly-dipeptides directly bind to the specific regions of Kapβ2.

**Figure 3.**
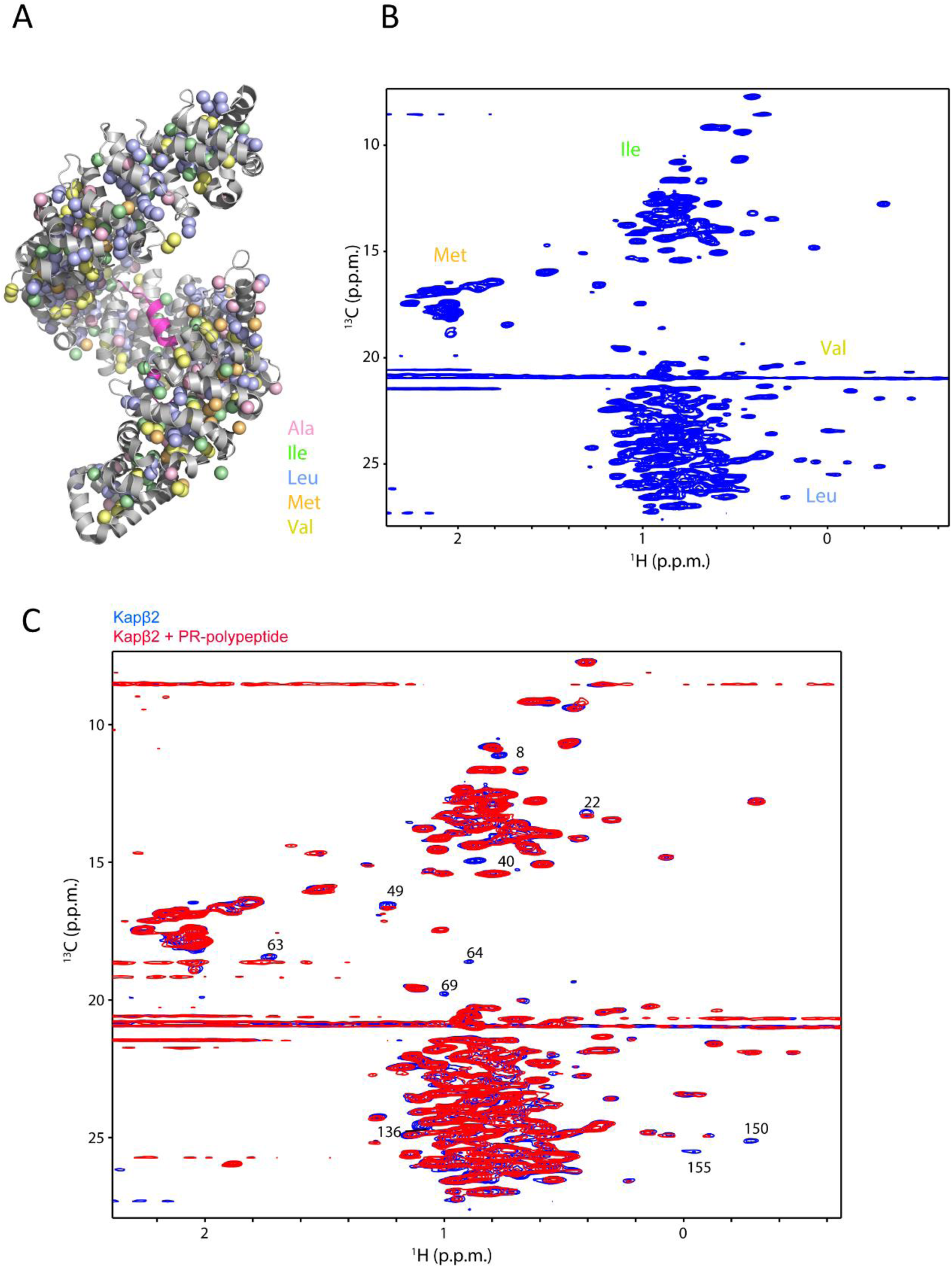
Interaction between Kapβ2 and PR poly-dipeptides investigated by solution NMR. (A) The crystal structure of Kapβ2 in complex with FUS PY-NLS (PDB ID: 4FDD). FUS PY-NLS is shown as magenta ribbon and Kapβ2 is show as gray ribbon with spheres representing methyl groups of Ala (pink), Ile (green), Leu (blue), Met (orange), and Val (yellow). Kapβ2 is enriched in the methyl-baring residues, indicating that sufficient information can be collected from the ^1^H-^13^C-correlated methyl NMR spectra. (B) ^1^H-^13^C-correlated methyl NMR spectra of [U-^2^ H; Ile-1-^13^CH_3_; Leu,Val-^13^CH_3_/C^2^H_3_]-labeled Kapβ2. (C) ^1^H-^13^C-correlated methyl NMR spectra of [U-^2^H; Ile-1-^13^CH_3_; Leu,Val-^13^CH_3_/C^2^H_3_]-labeled Kapβ2 in the absence (blue) and presence (red) of the PR poly-dipeptides. Significant perturbations were observed for several resonances. The significant representative perturbations are indicated by peak numbers.

### Interaction between Kapβ2 and PR20/M9M/FUS(1-500) investigated by solution NMR

In order to see if the perturbed resonances by the addition of PR20 are derived from the NLS-binding site of Kapβ2, we also performed NMR measurement for Kapβ2 in complex with M9M peptide; it has been shown to bind to NLS-binding site of Kapβ2 (24). We found that the binding of the M9M peptide induced significant perturbations to the many of the resonances of Kapβ2 (Fig. 4 and Fig. S2A). The significant perturbations suggest tight binding of the M9M peptide to Kapβ2, which is consistent with the high affinity reported in the previous study (24). Interestingly, several of the perturbed resonances by the addition of the PR poly-dipeptide were also caused by the binding of the M9M peptide (Fig. 3C and 4, Fig. S2A). Peaks overlapped between PR20 and M9M, indicating that the binding site for PR poly-dipeptide partially overlaps with the binding site for M9M peptide. Less significant perturbations by the addition of the PR-peptide can be explained by lower binding affinity for Kapβ2 (Fig. 2C, Table 1, Table S1).

**Figure 4.**
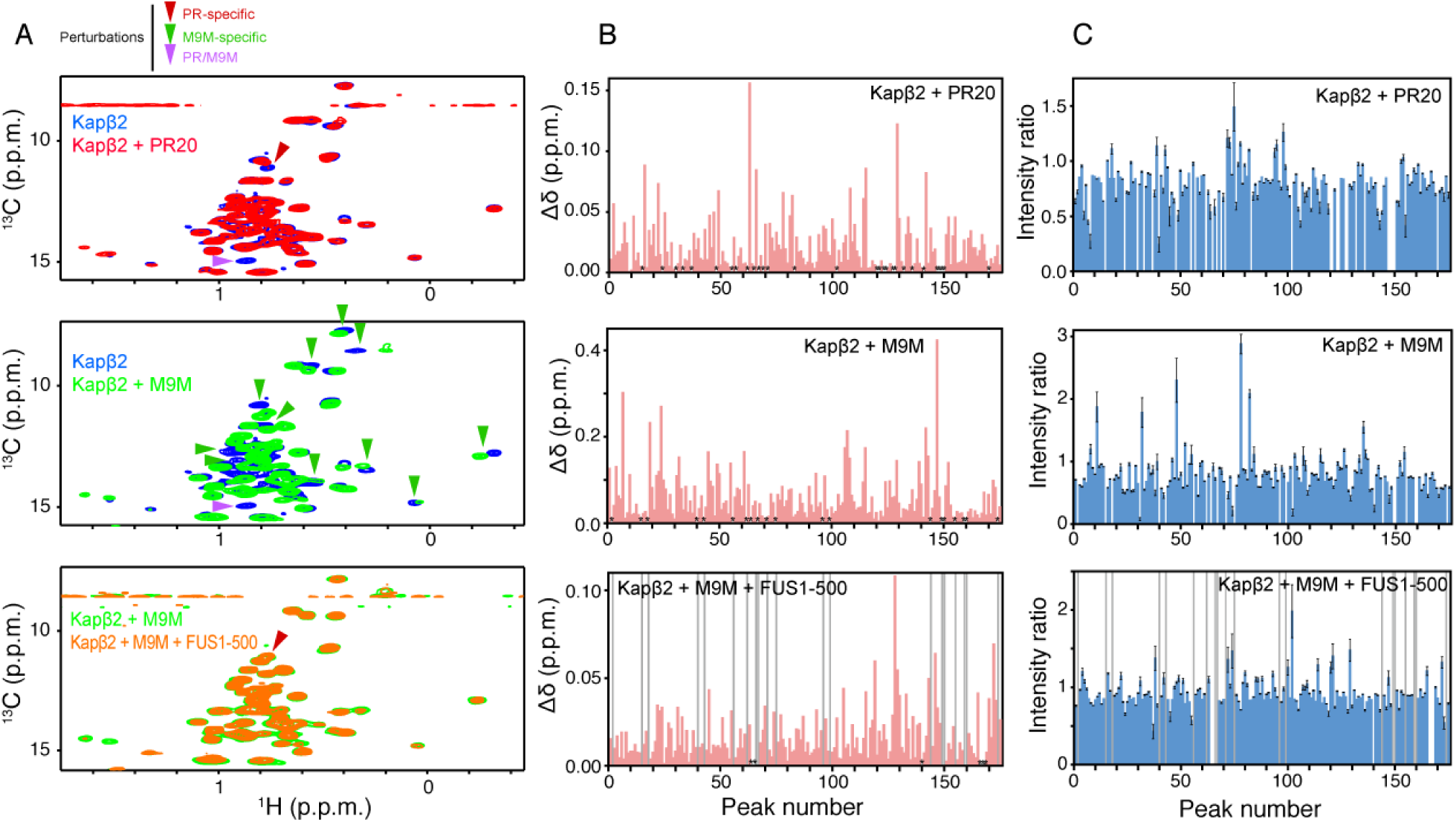
Interaction between Kapβ2 and PR20/M9M/FUS(1-500) investigated by solution NMR. (A) ^1^H-^13^C-correlated methyl NMR spectra of [U-^2^H; Ile-1-^13^CH_3_; Leu,Val-^13^CH_3_/C^2^H_3_]-labeled Kapβ2 in the absence (blue) and presence (red) of the PR poly-dipeptides. (top panel), in the absence (blue) and presence (green) of the M9M (middle panel), or in the presence of the M9M (green), and the presence of M9M and FUS(1-500) (orange) (bottom panel). The significant representative perturbations are indicated by arrow heads. The perturbations that are only seen for PR20 are indicated by red arrow heads. The perturbations that are only seen for M9M are indicated by green arrow heads. The perturbations that are common to PR and M9M are indicated by the purple arrow heads. For clarity, only the region of the Ile methyl resonances is shown. The full-range spectra are shown in Figure 3C (for top panel) and Figure S2 (for middle and bottom panels). Chemical shift difference (B) and intensity ratio (C) between the methyl resonances of Kapβ2 by the interaction with the PR poly-dipeptides (top panel), the M9M (middle panel), and FUS(1-500) (bottom panel). In panel (B), the resonances disappeared by the addition of the binding partner are indicated by the asterisks. In the bottom panels of (B) and (C), the resonances that are disappeared by complex formation with M9M are indicated by gray bars.

Perturbed resonances by the binding of M9M are expected to be derived around the NLS-binding site, as seen in the crystal structure of Kapβ2 in complex with PY-NLS (PDB code: 4FDD) (25). Here, we focused on the resonances that were perturbed both by the binding of PR20 and M9M (Fig. 3C and Fig. 4, Fig. S2A). The most significant perturbations were seen for the resonances numbered 40 (Ile), 49 (Met), 64 (Val), 150 (Leu), and 155 (Leu). The amino acid types were deduced from the chemical shift statistics of the methyl resonances (Fig. 3B). Our NMR data indicate that Ile, Leu, Val, and Met residues are involved in the interactions with PR20 and M9M. Given the fact that M9M binds to the PY-NLS-binding site of Kapβ2, these perturbed resonances can be attributed to the Kapβ2 residues close to PY-NLS as seen in the crystal structure of Kapβ2 in complex with PY-NLS (PDB code: 4FDD). These include the isoleucine residues (I457, I540, I642, I722, I773, I804), the leucine residues (L419, L539, L767), the valine residues (V643, V724), and the methionine residue (M308) (Fig. 5A, B). Among them, L539, I540, I642, V643 were found to be located at the negatively charged cavity of Kapβ2 (Fig. 5A). Taking into account the positive charge of the PR poly-dipeptide, these four residues located at the negatively charged cavity of Kapβ2 may be important for its interaction with PR poly-dipeptide. Strikingly, the molecular dynamics (MD) calculation demonstrated that PR poly-dipeptides are recognized by the negatively charged cavity and L419, I457, L539, I540, I642, V643, I722, are shown to be located in the binding site, which is highly consistent with the NMR data (Fig. 5C). Thus, NMR experiments corroborated by MD simulation indicated that the PR poly-dipeptide partially binds to the PY-NLS-binding site on Kapβ2.

**Figure 5.**
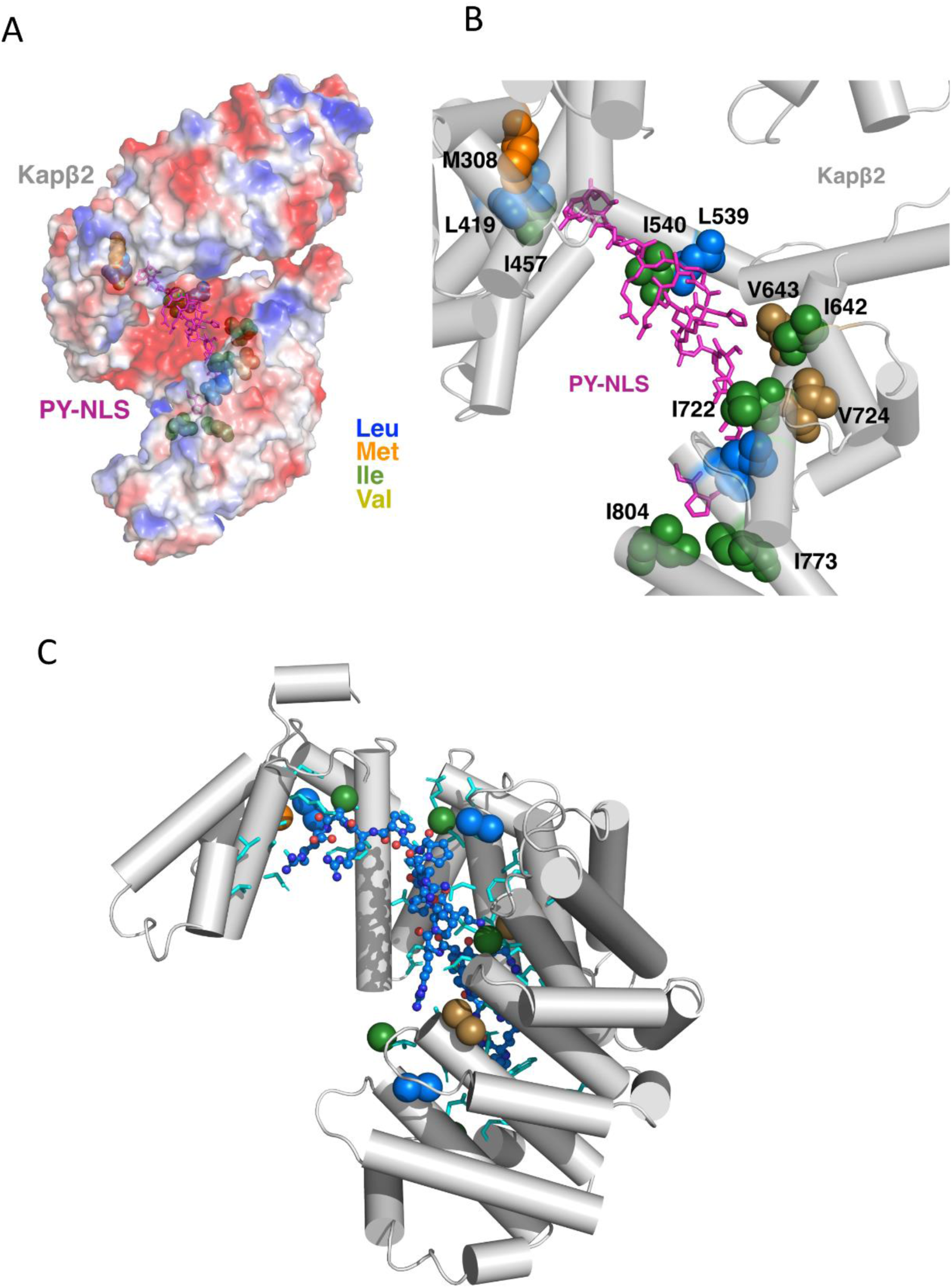
Structural model of PR interaction to Kapβ2. (A) Electrostatic potential of Kapβ2 (PDB code: 4FDD). PY-NLS colored magenta. Blue and red color shows positive charged and negative charged region, respectively. (B) Residues that are located on negative charged cavity. Leucine, isoleucine, Methionine and Valine are colored green, blue, orange and yellow, respectively. (C) MD calculated model of PR poly-dipeptides inside of Kapβ2 cavity.

A previous study demonstrated that Kapβ2 recognizes FUS regions other than NLS (16). In order to investigate the additional binding sites on Kapβ2 for FUS delta-NLS, we used NMR to investigate the interaction between Kapβ2 and FUS-ΔNLS(FUS1-500) in the presence of M9M peptide. Given that M9M peptide occupies the NLS-binding site on Kapβ2 (24), our data indicates that Kapβ2 recognizes FUS(1-500) using the sites other than NLS-binding site. The perturbed resonances induced by the addition of FUS are distinct from those perturbed by the binding of PR poly-dipeptide, indicating that the binding site for FUS(1-500) does not overlap with that for PR poly-dipeptide.

## Discussion

Neurodegenerative diseases are often linked to complications in nucleocytoplasmic transport (26,27). Nuclear pore complexes are composed of outer ring scaffold and centrally located LC-domains enriched with phenylalanine and glycine, generally called FG-domains that form permeability barrier between nucleus and cytoplasm. FG-domains form labile cross-β polymers and are the targets of toxic PR poly-dipeptides derived from *C9orf72* repeat expansions (3). Macro molecules go through nuclear pores upon exit and into the cellular nucleus. PR poly-dipeptides clog nuclear pores by binding to FG-domains and stabilizing FG-domains as polymeric, leading to nucleocytoplasmic transport defect. However, this does not fully account for the results from the genetic studies (19–21).

In this study, we show that PR poly-dipeptides disrupt chaperone activity of nuclear transport receptor Kapβ2 to FUS. We investigated the interactions between the PR poly-dipeptides and Kapβ2. In vitro binding assay showed that Kapβ2 exclusively binds to extended PR (PR18) poly-dipeptide. This result agrees with the toxicity of *C9orf72* expansions reported in familial ALS. Results from ITC experiments provided support for the inhibition of Kapβ2 by extended PR poly-dipeptide. The binding constant of PR18 and Kapβ2 was two folds stronger than that of full-length FUS and Kapβ2. Binding partners of Kapβ2 share PY-NLS and RGG domains (24), but detailed mechanisms driving Kapβ2 recognition of cargos remain to be elucidated.

Despite large size of Kapβ2 (100 kDa) for solution NMR, the use of the advanced NMR techniques including methyl-selective isotope labeling as well as methyl-transverse relaxation-optimized spectroscopy (TROSY) (22,23) achieved high-quality NMR spectra. This is one of the largest class of the proteins that are analyzed by high-resolution solution NMR. A series of the interaction analyses based on high-resolution NMR spectra have revealed that the PR poly-dipeptide binds to Kapβ2.

The binding sites on Kapβ2 partially overlaps with that used for the recognition of PY-NLS interactions, but does not overlap with that used for the recognition of FUS-ΔNLS (Fig. 3 and 4). Previous studies have shown that PY-NLS of FUS binds to the large center cavity of Kapβ2 that has negatively-charged regions (16,24). Negatively charged surface of the cavity may be targeted by arginines of PR poly-dipeptides. The binding of PR poly-dipeptide to the negatively-charged center cavity of Kapβ2 are further corroborated by MD calculation (Fig. 4). The amino acids within 5Å around the PR poly-dipeptide suggested by MD calculation were in good agreement with those indicated by NMR experiments. These data imply the possible mechanism of the toxicity of the PR poly-dipeptide toward Kapβ2-mediated nuclear transport, in which the PR poly-dipeptide interferes with the interaction between Kapβ2 and FUS NLS.

The current study revealed that PR poly-dipeptides bind nuclear import receptor Kapβ2 through PY-NLS recognition motif and compromise the ability of Kapβ2 to bind cargo proteins. This finding is an additional evidence how PR poly-dipeptides compromise protein-protein interaction within the cell. In addition to the ability of PR poly-dipeptides to bind and stabilize LC-polymers, the binding of PR poly-dipeptides to Kapβ2 explains why PR poly-dipeptides disrupt Kapβ2 to melt liquid-like droplets of FUS. The findings in this study offers mechanistic insights as to how *C9orf72* repeat expansion disables nucleocytoplasmic transport and causes neurodegenerative diseases.

## Experimental procedures

### Constructs, protein expression and purification

Kapβ2 was expressed from GST-fusion constructs using pGEX6P-1 vector. FUS and PRn proteins were expressed from MBP-fusion constructs using the pMAL-TEV vector. All recombinant proteins were expressed individually in BL21(DE3) *E.coli* cells induced with 0.5 mM ispropyl-β-d-1-thiogrlactoside (IPTG) for 12 hours at 20°C. Bacteria expressing Kapβ2 was lysed with sonicator in buffer containing 50 mM Tris pH 7.5, 200 mM NaCl, 20%(v/v) glycerol, 2 mM DTT. Kapβ2 was purified using GSH Spharose beads (GS4B, GE Healthcare), cleaved with HRV3C protease, anion exchange chromatography (HiTrap Q HP, GE Healthcare) and gel filtration chromatography (Superdex200 16/60, GE Healthcare) in buffer containing 20 mM HEPES pH 7.4, 150 mM NaCl, 2 mM DTT, 2 mM Mg(OAc)_2_ and 10% glycerol. MBP-FUS and MBP-PRn proteins were lysed in 50 mM Tris pH7.5, 1.5 M NaCl, 10% glycerol, 2 mM DTT. MBP-fusion proteins were purified by affinity chromatography using amylose resin eluted with buffer containing 50 mM Tris pH7.5, 150 mM NaCl, 10% glycerol, 2 mM DTT and 20 mM Maltose. It was then further purified for MBP-PRn by cation-exchange chromatography (HiTrap SP HP, GE Healthcare) and gel filtration chromatography (Superdex200 16/60, GE Healthcare).

### Turbidity assay and Imaging of turbid solution

Prior to adding TEV protease, 8 μM MBP-FUS, ± 8 μM Kapβ2 ±MBP-PR18 were mixed buffer containing 20 mM Hepes pH7.4, 150 mM NaCl, 10% Glycerol, 2 mM Mg(Oac)_2_, 20 μM Zn(Oac)_2_ and 2 mM DTT to reaction volumes of 100 μL. TEV protease was added to the premixture, to a final concentration of 40 μg/mL then incubated at 30°C for 60 min to digest all MBP-fusion protein. The solution let cool down to 20°C then measure OD 395 nm using plate reader. For the imaging experiment, turbid solutions were imaged with a fluorescence microscopy (Olympus BX53).

### Hydrogel binding assay

GFP and mCherry fusion LC domain of FUS (residue 2-214) and hnRNPA2 (residue 181-341) were expressed in *E.coli* BL21(DE3) cells with 0.5 mM IPTG at 20°C for overnight and purified as previously described (1). Expression plasmids for recombinant proteins were obtained from Steven L. McKnight Laboratory. Hydrogel droplets of mCherry:LC domain of FUS and hnRNPA2 were prepared as previously reported (1). For hydrogel binding assays, purified GFP-fused proteins were diluted to 1 µM in the buffer (20mM Tris-HCl pH 7.5, 150mM NaCl, 20mM BME, 0.1mM PMSF, 0.5mM EDTA) and pipetted into the hydrogel dish. After overnight incubation, horizontal sections of the droplets were scanned with excitation wavelengths on a confocal microscope (FLUOVIEW FV3000, OLYMPUS). PR20-HA peptides were synthesized by SCRUM Inc. (Japan).

### Pull-down binding assay

In-vitro pull-down binding assays were performed using GST-PR18 or GST-Kapββ2 immobilized on GSH sepharose beads (GE Healthcare). Four μg GST proteins were immobilized on beads. Thirty μL of GST-proteins beads are incubated with equal molar Kapβ2, MBP-PRn for 20-30 min and washed three times with buffer containing 20mM Hepes pH7.4, 150 mM NaCl, 10% Glycerol, 2mM Mg(OAc)_2_ and 2mM DTT. Bound proteins were separated by SDS-PAGE and stained with Coomassie Brilliant Blue.

### Isothermal titration calorimetry (ITC)

ITC experiments were performed with a Malvern iTC200 calorimeter (Malvern Instruments). Proteins were dialyzed overnight against buffer containing 20mM Hepes pH7.4, 150mM NaCl, 10% Glycerol and 2mM β-mercaptoethanol. 200μM MBP-PR18 was titrated into the sample cell containing 20μM Kapβ2(ΔLoop, residues 321-371 were replaced with GGSGGSGS linker). ITC experiments were performed at 25°C with 19 rounds of 2μL injections. Data was analyzed using Origin software.

### Expression and purification of isotopically labeled Kapβ2

The Kapβ2 expression plasmid was transformed into BL21(DE3) cells. The protein sample with ^1^H,^13^C-labeled methyl groups in deuterium background were prepared as described previously (22). The cells were grown in medium with ^15^NH_4_Cl (2 gL^−1^) (CIL) and ^2^H_7_-glucose (2 gL^−1^) (CIL) in 99.9% ^2^H_2_O (Isotec). The precursors for methyl groups of Ile, Val, and Leu, α-ketobutyric acid (50 mg L^−1^) and α-ketoisovaleric acid (80 mg L^−1^), and [^13^CH_3_] methionine (50 mg ^L−1^) were added to the culture one hour before the addition of IPTG. Protein expression was induced by the addition of 0.5 mM isopropyl-β-D-1-thiogalactopyranoside (IPTG) at OD_600_ ~0.6, followed by ~16 hours of incubation at 25°C. Cells were harvested and resuspended in the lysis buffer containing 50mM HEPES pH 7.4, 150mM NaCl, 20 % Glycerol, 2mM DTT, 2mM EDTA. Cells were disrupted by sonicator and centrifuged at 18,000rpm for 45 min. Proteins were purified using GS4B resin (GE healthcare) and eluted with the buffer containing 20mM HEPES pH7.4, 20mM NaCl, 2 mM EDTA, 10% Glycerol, 30mM GSH. The GST tag was removed by HRV 3C protease at 4°C for ~16 h. The protein was further purified by anion exchange using HiTrap Q FF (GE healthcare) with the buffer containing 20mM Imidazole pH 6.5, 20-1000mM NaCl, 2mM EDTA, 2mM DTT, 20% Glycerol, followed by gel filtration using Superdex 200 16/60 (GE Healthcare) with the buffer containing 20mM HEPES pH7.4,150mM NaCl, 2mM MgCl_2_, 2mM DTT, 2% Glycerol. Protein concentration was determined spectrophotometrically at 280 nm using the corresponding extinction coefficient.

### NMR experiments

The isotopically labeled Kapβ2 was prepared in the NMR buffer containing 20 mM ^2^H-Tris pH 7.4, 150 mM NaCl, 2 mM MgCl_2_, 2mM DTT, 2% glycerol, and concentrated to 20 μM. NMR experiments were performed on Bruker 600 MHz NMR at 20°C. Perturbation of the side chain methyl resonances were monitored using ^1^H-^13^C heteronuclear multiple quantum coherence (HMQC), respectively. Spectra were processed using the NMRPipe software (28). Data analysis was performed by Olivia software (http://fermi.pharm.hokudai.ac.jp/olivia/). The perturbations of the resonances were evaluated by chemical shift change or intensity change. Chemical shift perturbations (CSPs) for the methyl groups were calculated by the equation;

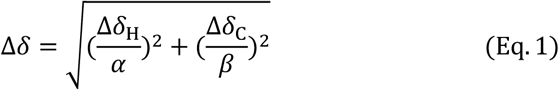

where ∆*δ*_H_ and ∆*δ*_C_ are the chemical shift changes of ^1^H and ^13^C, respectively, by the addition of the ligand, and α and β are chemical shift distribution of ^1^H and ^13^C, respectively, of methyl groups as reported in the Biological Resonance Data Bank (http://www.bmrb.wisc.edu) (29). Errors for the intensity ratio were estimated based on the peak intensity and noise level.

In order to investigate the interaction between Kapβ2 and PR poly-dipeptide by NMR, the NMR spectra of the isotopically labeled Kapβ2 in the absence and presence of the synthetic peptide of (Pro-Arg)_20_-HA were measured. In order to investigate the interaction between Kapβ2 and the M9M peptide, the Kapβ2-M9M complex was prepared as follows: The purified GST-M9M protein was added to the isotopically labeled Kapβ2, followed by the removal of the GST tag by TEV protease at RT for 2h and then 4°C for ~16 h before it was buffer exchanged into the NMR buffer using Amicon Ultra-4 50k (Merck). In order to investigate the interaction between Kapβ2 and FUS(1-500) by NMR, the NMR spectra of the isotopically labeled Kapβ2 in complex with the M9M peptide in the absence and presence of FUS(1-500) were measured.

### MD simulation

Molecular dynamics (MD) simulations were performed by Maestro (Schrödinger) with Desmond program (D. E. Shaw Research). The input structure for MD simulations were prepared by taking chain A of the crystallographic dimer from the crystal structure of Kapβ2 in complex with FUS (PDB: 5YVG), and mutating side chains of chain X to PR. The protein and PR poly-dipeptide were solvated with a rectangular box of TIP3P water molecules and neutralized by adding sodium ions. The simulation system was equilibrated with the default protocol of the Desmond program. A 15ns MD simulation was performed under the following conditions: equilibration of simulation was performed using the isothermal–isobaric ensemble (NPT), the van der Waals interaction between atoms separated by over 10Å were cut off.

## Acknowledgements

The authors thank Steven L. McKnight (UT Southwestern Medical Center), Yuh-Min Chook (UT Southwestern Medical Center), Masato Kato (UT Southwestern Medical Center), Kentaro Shiraki (University of Tsukuba), Yoichi Shinkai (National Institute of Advanced Industrial Science and Technology), Tatsuya Niwa (Tokyo Institute of Technology), Hideki Taguchi (Tokyo Institute of Technology) and Ms. Keren-Happuch E Fan Fen for their critical reading of the manuscript. The authors thank Fumio Takahashi for his help with microscopic imaging and Yoichi Takeda for his help with ITC data collection. The authors thank Hiroyuki Kumeta for his help in setting up NMR experiments. The NMR experiments were performed at Hokkaido University Advanced NMR Facility, a member of NMR Platform.

## FOOTNOTES

This work was supported by grants from AMED Brain/MINDS Beyond [JP19dm0307032] to E.M., JSPS KAKENHI [JP17H07031 to E.M., JP18H06202 to H.N., JP19H04945, JP19K06504, JP18H05229, JP17H05657, JP17H05867 to T.S., JP19H05769 to K.I., JP19K16060 to T.Y], Takeda Science Foundation to E.M. and T.S., Kanzawa Medical Research Foundation to E.M., Uehara Memorial Foundation to E.M., Nakatomi Foundation to E.M., Konica Minolta Science and Technology Foundation to E.M., Naito Foundation to E.M., MSD Life Science Foundation to E.M., Mochida Memorial Foundation for Medical and Pharmaceutical Research to E.M., SENSHIN Medical Research Foundation to E.M., Terumo Foundation for Life Sciences and Arts to E.M., Nara Kidney Disease Research Foundation to E.M., Novartis Research Grants to E.M., H.N. and K.S., Nara Medical University Grant-in-Aid for Collaborative Research Projects to K.S., Akiyama Life Science Foundation Grants-in-Aid to T.S., Northern Advancement Center for Science and Technology Grants-in-Aid to T.S., Daiichi Sankyo Foundation of Life Science Grants-in-Aid to T.Y., The Nakabayashi Trust For ALS Research Grants-in-Aid to T.Y., Integrated Research Consortium on Chemical Sciences to Y.A., Izumi Science and Technology Foundation to Y.A., and by unrestricted funds provided to E.M. from Dr. Taichi Noda (KTX Corp., Aichi, Japan) and Dr. Yasuhiro Horii (Koseikai, Nara, Japan).

## The abbrevations used are

LC: low-complexity
ALS: amyotrophic lateral sclerosis
NLS: nuclear localization signal
PY-NLS: proline-tyrosine nuclear localization signal
FUS: fused in sarcoma
hnRNP: heterogeneous nuclear ribonucleo-protein
LC polymer: labile cross-β polymer formed from LC-domains
MSP: multisystem proteinopathy
FTD: frontotemporal dementia
PR: proline:arginine
GR: glycine:arginine
FG: phenylalanine:glycine
MBP: maltose binding protein
ITC: isothermal titration calorimetry
NMR: nuclear magnetic resonance
TROSY: methyl-transverse relaxation-optimized spectroscopy
MD: molecular dynamics

## References

1. Kato, M., Han, T. W., Xie, S., Shi, K., Du, X., Wu, L. C., Mirzaei, H., Goldsmith, E. J., Longgood, J., Pei, J., Grishin, N. V., Frantz, D. E., Schneider, J. W., Chen, S., Li, L., Sawaya, M. R., Eisenberg, D., Tycko, R., and McKnight, S. L. (2012) Cell-free formation of RNA granules: low complexity sequence domains form dynamic fibers within hydrogels. Cell 149, 753–767

2. Lin, Y., Mori, E., Kato, M., Xiang, S., Wu, L., Kwon, I., and McKnight, S. L. (2016) Toxic PR Poly-Dipeptides Encoded by the C9orf72 Repeat Expansion Target LC Domain Polymers. Cell 167, 789–802.e712

3. Shi, K. Y., Mori, E., Nizami, Z. F., Lin, Y., Kato, M., Xiang, S., Wu, L. C., Ding, M., Yu, Y., Gall, J. G., and McKnight, S. L. (2017) Toxic PRn poly-dipeptides encoded by the C9orf72 repeat expansion block nuclear import and export. Proc Natl Acad Sci U S A 114, E1111–e1117

4. Xiang, S., Kato, M., Wu, L. C., Lin, Y., Ding, M., Zhang, Y., Yu, Y., and McKnight, S. L. (2015) The LC Domain of hnRNPA2 Adopts Similar Conformations in Hydrogel Polymers, Liquid-like Droplets, and Nuclei. Cell 163, 829–839

5. Han, T. W., Kato, M., Xie, S., Wu, L. C., Mirzaei, H., Pei, J., Chen, M., Xie, Y., Allen, J., Xiao, G., and McKnight, S. L. (2012) Cell-free formation of RNA granules: bound RNAs identify features and components of cellular assemblies. Cell 149, 768–779

6. Kwon, I., Kato, M., Xiang, S., Wu, L., Theodoropoulos, P., Mirzaei, H., Han, T., Xie, S., Corden, J. L., and McKnight, S. L. (2013) Phosphorylation-regulated binding of RNA polymerase II to fibrous polymers of low-complexity domains. Cell 155, 1049–1060

7. Murray, D. T., Kato, M., Lin, Y., Thurber, K. R., Hung, I., McKnight, S. L., and Tycko, R. (2017) Structure of FUS Protein Fibrils and Its Relevance to Self-Assembly and Phase Separation of Low-Complexity Domains. Cell 171, 615–627.e616

8. Kato, M., and McKnight, S. L. (2018) A Solid-State Conceptualization of Information Transfer from Gene to Message to Protein. Annu Rev Biochem 87, 351–390

9. Murray, D. T., Zhou, X., Kato, M., Xiang, S., Tycko, R., and McKnight, S. L. (2018) Structural characterization of the D290V mutation site in hnRNPA2 low-complexity-domain polymers. Proc Natl Acad Sci U S A 115, E9782–e9791

10. Kwon, I., Xiang, S., Kato, M., Wu, L., Theodoropoulos, P., Wang, T., Kim, J., Yun, J., Xie, Y., and McKnight, S. L. (2014) Poly-dipeptides encoded by the C9orf72 repeats bind nucleoli, impede RNA biogenesis, and kill cells. Science 345, 1139–1145

11. Mizielinska, S., Gronke, S., Niccoli, T., Ridler, C. E., Clayton, E. L., Devoy, A., Moens, T., Norona, F. E., Woollacott, I. O. C., Pietrzyk, J., Cleverley, K., Nicoll, A. J., Pickering-Brown, S., Dols, J., Cabecinha, M., Hendrich, O., Fratta, P., Fisher, E. M. C., Partridge, L., and Isaacs, A. M. (2014) C9orf72 repeat expansions cause neurodegeneration in Drosophila through arginine-rich proteins. Science 345, 1192–1194

12. Lee, K.-H., Zhang, P., Hong, Mitrea, D. M., Sarkar, M., Freibaum, B. D., Cika, J., Coughlin, M., Messing, J., Molliex, A., Maxwell, B. A., Nam, Temirov, J., Moore, J., Kolaitis, R.-M., Shaw, T. I., Bai, B., Peng, J., Kriwacki, R. W., and J. (2016) C9orf72 Dipeptide Repeats Impair the Assembly, Dynamics, and Function of Membrane-Less Organelles. 167, 774–788.e717

13. Kato, M., Yang, Y. S., Sutter, B. M., Wang, Y., McKnight, S. L., and Tu, B. P. (2019) Redox State Controls Phase Separation of the Yeast Ataxin-2 Protein via Reversible Oxidation of Its Methionine-Rich Low-Complexity Domain. Cell 177, 711–721.e718

14. Hofweber, M., Hutten, S., Bourgeois, B., Spreitzer, E., Niedner-Boblenz, A., Schifferer, M., Ruepp, M. D., Simons, M., Niessing, D., Madl, T., and Dormann, D. (2018) Phase Separation of FUS Is Suppressed by Its Nuclear Import Receptor and Arginine Methylation. Cell 173, 706–719.e713

15. Yang, Y. S., Kato, M., Wu, X., Litsios, A., Sutter, B. M., Wang, Y., Hsu, C. H., Wood, N. E., Lemoff, A., Mirzaei, H., Heinemann, M., and Tu, B. P. (2019) Yeast Ataxin-2 Forms an Intracellular Condensate Required for the Inhibition of TORC1 Signaling during Respiratory Growth. Cell 177, 697–710.e617

16. Yoshizawa, T., Ali, R., Jiou, J., Fung, H. Y. J., Burke, K. A., Kim, S. J., Lin, Y., Peeples, W. B., Saltzberg, D., Soniat, M., Baumhardt, J. M., Oldenbourg, R., Sali, A., Fawzi, N. L., Rosen, M. K., and Chook, Y. M. (2018) Nuclear Import Receptor Inhibits Phase Separation of FUS through Binding to Multiple Sites. Cell 173, 693–705.e622

17. Guo, L., Kim, H. J., Wang, H., Monaghan, J., Freyermuth, F., Sung, J. C., O’Donovan, K., Fare, C. M., Diaz, Z., Singh, N., Zhang, Z. C., Coughlin, M., Sweeny, E. A., DeSantis, M. E., Jackrel, M. E., Rodell, C. B., Burdick, J. A., King, O. D., Gitler, A. D., Lagier-Tourenne, C., Pandey, U. B., Chook, Y. M., Taylor, J. P., and Shorter, J. (2018) Nuclear-Import Receptors Reverse Aberrant Phase Transitions of RNA-Binding Proteins with Prion-like Domains. Cell 173, 677–692.e620

18. Qamar, S., Wang, G., Randle, S. J., Ruggeri, F. S., Varela, J. A., Lin, J. Q., Phillips, E. C., Miyashita, A., Williams, D., Strohl, F., Meadows, W., Ferry, R., Dardov, V. J., Tartaglia, G. G., Farrer, L. A., Kaminski Schierle, G. S., Kaminski, C. F., Holt, C. E., Fraser, P. E., Schmitt-Ulms, G., Klenerman, D., Knowles, T., Vendruscolo, M., and St George-Hyslop, P. (2018) FUS Phase Separation Is Modulated by a Molecular Chaperone and Methylation of Arginine Cation-pi Interactions. Cell 173, 720–734.e715

19. Zhang, K., Donnelly, C. J., Haeusler, A. R., Grima, J. C., Machamer, J. B., Steinwald, P., Daley, E. L., Miller, S. J., Cunningham, K. M., Vidensky, S., Gupta, S., Thomas, M. A., Hong, I., Chiu, S. L., Huganir, R. L., Ostrow, L. W., Matunis, M. J., Wang, J., Sattler, R., Lloyd, T. E., and Rothstein, J. D. (2015) The C9orf72 repeat expansion disrupts nucleocytoplasmic transport. Nature 525, 56–61

20. Freibaum, B. D., Lu, Y., Lopez-Gonzalez, R., Kim, N. C., Almeida, S., Lee, K. H., Badders, N., Valentine, M., Miller, B. L., Wong, P. C., Petrucelli, L., Kim, H. J., Gao, F. B., and Taylor, J. P. (2015) GGGGCC repeat expansion in C9orf72 compromises nucleocytoplasmic transport. Nature 525, 129–133

21. Jovicic, A., Mertens, J., Boeynaems, S., Bogaert, E., Chai, N., Yamada, S. B., Paul, J. W., 3rd, Sun, S., Herdy, J. R., Bieri, G., Kramer, N. J., Gage, F. H., Van Den Bosch, L., Robberecht, W., and Gitler, A. D. (2015) Modifiers of C9orf72 dipeptide repeat toxicity connect nucleocytoplasmic transport defects to FTD/ALS. Nat Neurosci 18, 1226–1229

22. Saio, T., Guan, X., Rossi, P., Economou, A., and Kalodimos, C. G. (2014) Structural Basis for Protein Antiaggregation Activity of the Trigger Factor Chaperone. Science 344, 1250494–1250494

23. Saio, T., Kawagoe, S., Ishimori, K., and Kalodimos, C. G. (2018) Oligomerization of a molecular chaperone modulates its activity. eLife 7

24. Cansizoglu, A. E., Lee, B. J., Zhang, Z. C., Fontoura, B. M. A., and Chook, Y. M. (2007) Structure-based design of a pathway-specific nuclear import inhibitor. Nature Structural & Molecular Biology 14, 452–454

25. Zhang, Z. C., and Chook, Y. M. (2012) Structural and energetic basis of ALS-causing mutations in the atypical proline–tyrosine nuclear localization signal of the Fused in Sarcoma protein (FUS). Proceedings of the National Academy of Sciences 109, 12017–12021

26. Folkmann, A. W., Collier, S. E., Zhan, X., Aditi, Ohi, M. D., and Wente, S. R. (2013) Gle1 functions during mRNA export in an oligomeric complex that is altered in human disease. Cell 155, 582–593

27. Hirano, M., Furiya, Y., Asai, H., Yasui, A., and Ueno, S. (2006) ALADINI482Scauses selective failure of nuclear protein import and hypersensitivity to oxidative stress in triple A syndrome. Proceedings of the National Academy of Sciences 103, 2298–2303

28. Delaglio, F., Grzesiek, S., Vuister, G. W., Zhu, G., Pfeifer, J., and Bax, A. (1995) NMRPipe: a multidimensional spectral processing system based on UNIX pipes. J Biomol NMR 6, 277–293

29. Rosenzweig, R., Farber, P., Velyvis, A., Rennella, E., Latham, M. P., and Kay, L. E. (2015) ClpB N-terminal domain plays a regulatory role in protein disaggregation. Proceedings of the National Academy of Sciences 112, E6872–E6881

