## Supplemental Table and Figures for "Toxic PR poly-dipeptides encoded by the *C9orf72* repeat expansion target Kapβ2 and dysregulate phase separation of low-complexity domains"

**Table S1. Kd value against Kapβ2 determined by ITC**

| Ligand | Tag | Kd (nM) | Reference | Buffer |
|---|---|---|---|---|
| PR18 | MBP | 81.3 | this work | 20 mM HEPES pH7.4, 150 mM NaCl, 10% glycerol, 2 mM bME |
| FUS(476-526) | MBP | 71.5 | Yoshizawa et al., Cell, 2018 | 20 mM HEPES pH7.4, 150 mM NaCl, 10% glycerol, 2 mM bME |
| FUS(453-526) | MBP | 173 | Yoshizawa et al., Cell, 2018 | 20 mM HEPES pH7.4, 150 mM NaCl, 10% glycerol, 2 mM bME |
| FUS(371-526) | MBP | 209 | Yoshizawa et al., Cell, 2018 | 20 mM HEPES pH7.4, 150 mM NaCl, 10% glycerol, 2 mM bME |
| FUS(1-526) | MBP | 158.7 | Yoshizawa et al., Cell, 2018 | 20 mM HEPES pH7.4, 150 mM NaCl, 10% glycerol, 2 mM bME |
| M9NLS-A1(257-305) | MBP | 42 | Lee et al., Cell, 2006 | 20 mM Tris pH7.5, 100 mM NaCl, 2 mM bME |
| hnRNPM(41-70) | MBP | 10 | Cansizoglu et al., NSMB, 2007 | 20 mM Tris pH7.5, 100 mM NaCl, 2 mM bME |
| M9M | MBP | 0.107 | Cansizoglu et al., NSMB, 2007 | 20 mM Tris pH7.5, 100 mM NaCl, 2 mM bME |
| FUS(504-526) | - | 20000 | Dormann et al., EMBO J, 2012 | 20 mM Na-phosphase buffer pH6.8, 50 mM NaCl, 1 mM EDTA, 1 mM DTT |
| FUS(489-526) | - | 2800 | Dormann et al., EMBO J, 2012 | 20 mM Na-phosphase buffer pH6.8, 50 mM NaCl, 1 mM EDTA, 1 mM DTT |
| FUS(473-503) | - | 11700 | Dormann et al., EMBO J, 2012 | 20 mM Na-phosphase buffer pH6.8, 50 mM NaCl, 1 mM EDTA, 1 mM DTT |
| FUS(454-526) | - | 600 | Dormann et al., EMBO J, 2012 | 20 mM Na-phosphase buffer pH6.8, 50 mM NaCl, 1 mM EDTA, 1 mM DTT |

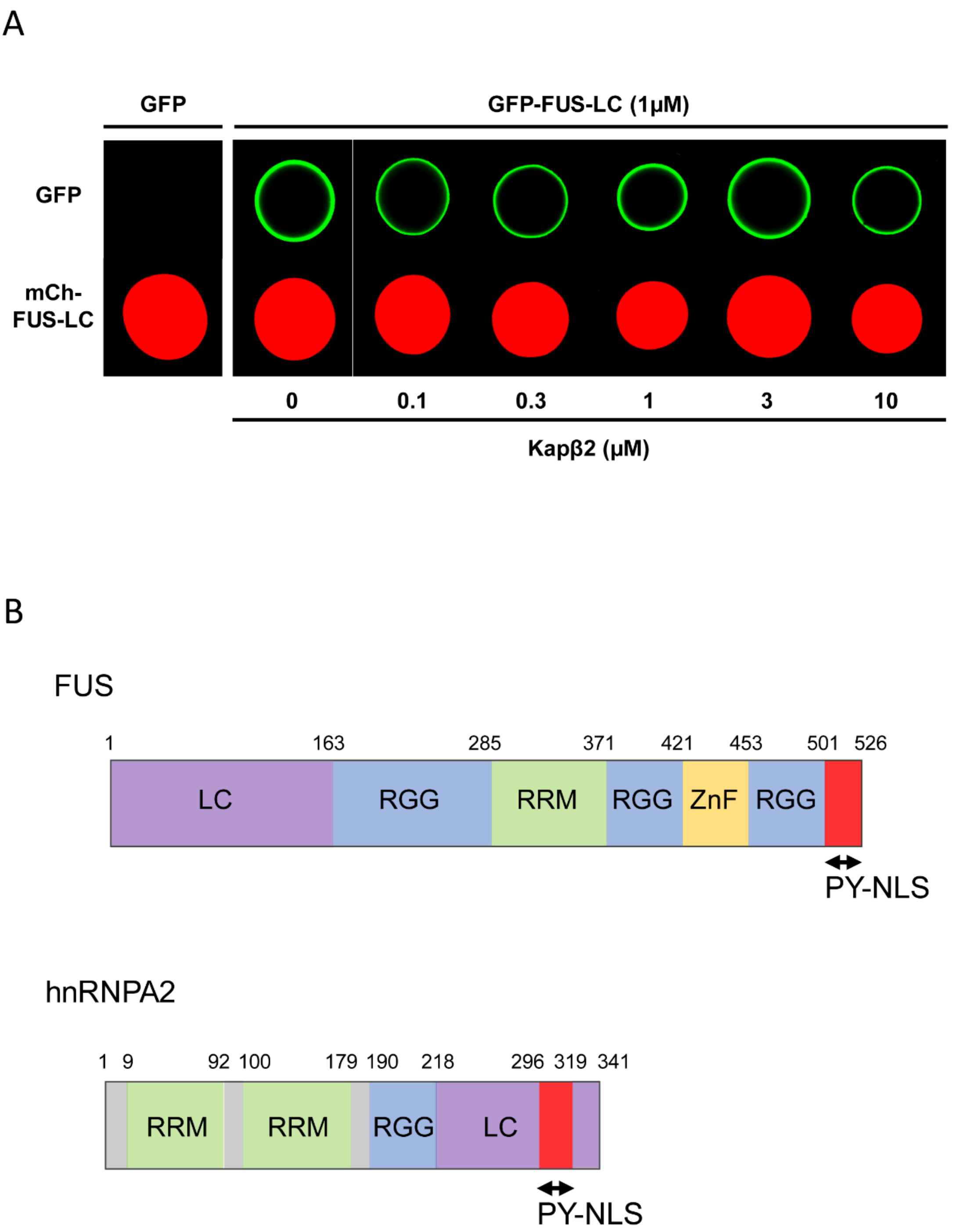


**Figure S1.** (A) Hydrogel droplets made from mCherry:LC domain of FUS (lower images) were incubated with 1μM of GFP (left panel) and GFP:FUS-LC (right panel) and visualized by confocal microscopy. GFP:FUS-LC was challenged for homotypic polymer extension in the presence of different concentration of Kapβ2 (left to right: 0.1μM, 0.3μM, 1μM, 3μM, 10μM, respectively). (B) Domain architecture of FUS and hnRNPA2. FUS has a PY-NLS in the C terminal. PY-NLS is located in the middle of LC domain of hnRNPA2.

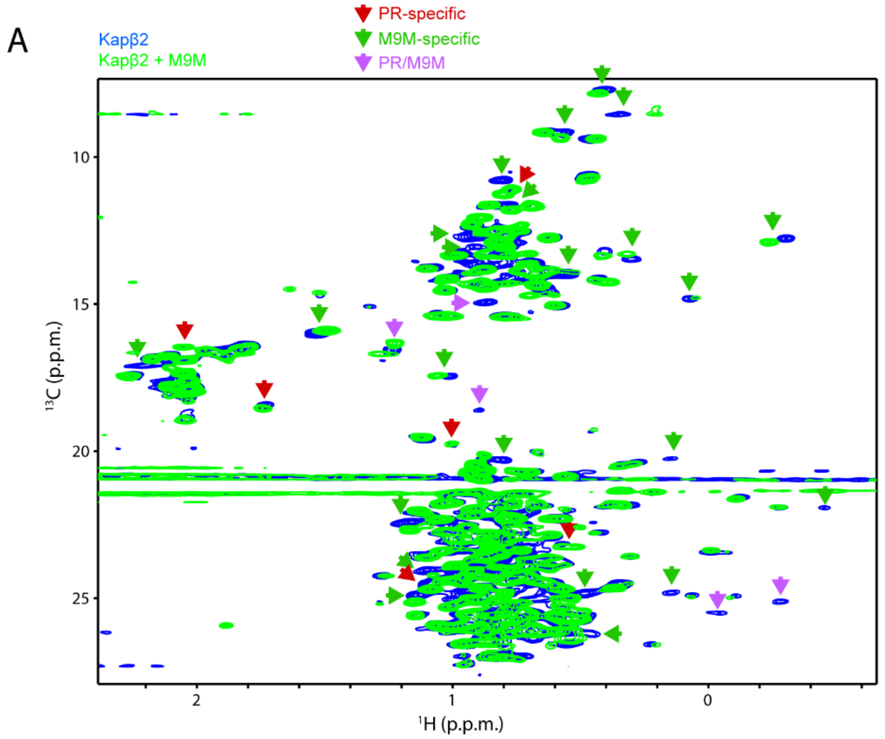


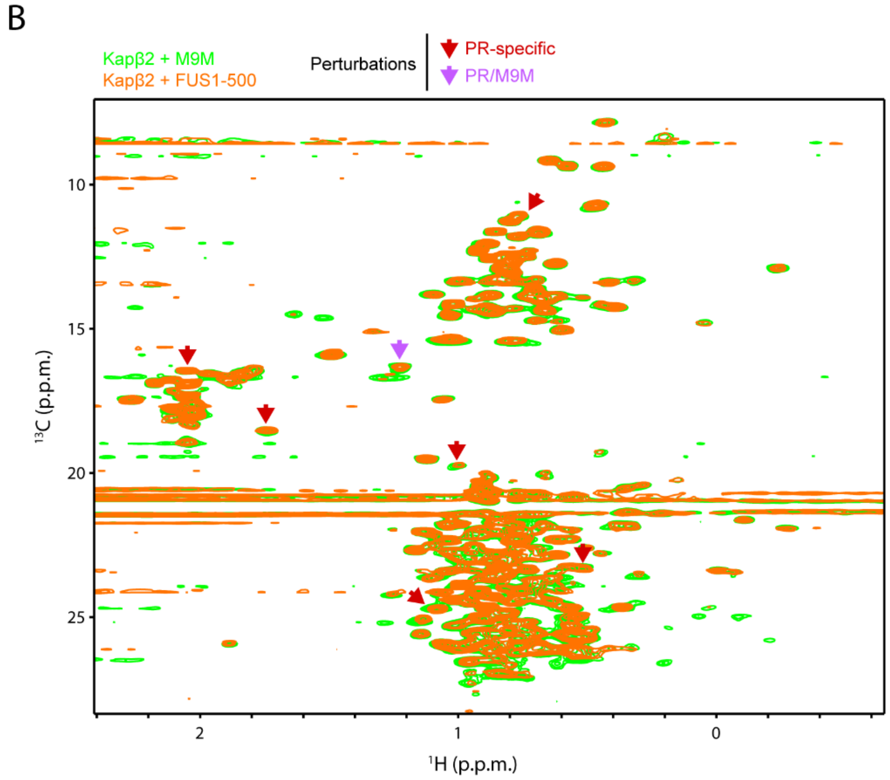


**Figure S2.** (A) $^{1}$H-$^{13}$C-correlated methyl NMR spectra of [U-$^{2}$H; Ile-1-$^{13}CH_3$; Leu,Val-$^{13}CH_3/C^2H_3$]-labeled Kapβ2 (blue) and that in complex with M9M (green). (B) $^{1}$H-$^{13}$C-correlated methyl NMR spectra of [U-$^{2}$H; Ile-1-$^{13}CH_3$; Leu,Val-$^{13}CH_3/C^2H_3$]-labeled Kapβ2 in complex with M9M, in the absence (green) and presence FUS(1-500) (orange). The significant representative perturbations are indicated by arrows. The perturbations that are only seen for PR20 are indicated by red arrows. The perturbations that are only seen for M9M are indicated by green arrows. The perturbations that are common to PR and M9M are indicated by the purple arrows.
